# Microstructural determinants of lens stiffness in rat versus guinea pig lenses

**DOI:** 10.1101/2021.02.15.431302

**Authors:** Justin Parreno, Kalekidan Abera, Sandeep Aryal, Karen E. Forbes, Velia M. Fowler

**Affiliations:** Department of Biological Sciences, University of Delaware; College of Agriculture and Natural Resources, University of Delaware

## Abstract

Proper ocular lens function requires lens biomechanical flexibility which is lost in presbyopia during aging. As increasing lens size has been shown previously to correlate with lens biomechanical stiffness in aging, we tested the hypothesis that whole lens size determines gross biomechanical stiffness. We used an allometric approach to evaluate this hypothesis by comparing lenses from three rodent species (mouse, rats and guinea pigs) of varying size. While rat lenses are larger and stiffer than mouse lenses, guinea pig lenses are even larger than rat lenses but are softer than the rat lens. This indicates that lens size is not a sole determinant of lens stiffness and disproves our hypothesis. Therefore, we investigated the scaling of lens microstructural features that could potentially explain the differences in biomechanical stiffness between rat and guinea pig lenses, including lens capsule thickness, epithelial cell area, fiber cell widths, suture organization, and nuclear size. Capsule thickness, epithelial cell area, and fiber cell widths scaled with lens size (i.e., greater in guinea pig lenses than rats), indicating that sizes of these features do not correlate with the stiffness of rat lenses, while suture organization was similar between rats and guinea pigs. However, we found that the hard rat lens nucleus occupies a greater fraction of the lens than the guinea pig lens nucleus, suggesting a role for nuclear size in determining whole lens stiffness. Therefore, while many features contribute to lens biomechanical properties, the size of the lens nucleus with respect to the size of the lens could be a major determinant of lens stiffness in rats versus guinea pigs.

## INTRODUCTION

The ocular lens is a clear transparent tissue that is responsible for fine focusing light onto the retina. In humans and primates, the lens changes shape in the process of accommodation and disaccommodation to generate a clear image onto the retina for near and far vision, respectively. This lens function is strongly linked to its biomechanical properties. A major age-related pathology of the lens is presbyopia or far-sightedness where the ability for lens shape change is hindered (Glasser and Campbell, 1999; Heys et al., 2004; Weeber et al., 2005). This inability for shape change has been associated with increase in lens biomechanical stiffness during aging, which has been demonstrated in humans (Burd et al., 2011; Fisher, 1971; Glasser and Campbell, 1998; Heys et al., 2004; Krueger et al., 2001; Pau and Kranz, 1991; Weeber and van der Heijde, 2007) and rodent animal models (Baradia et al., 2010; Cheng et al., 2016; Gokhin et al., 2012; Schindlery, 1989).

The lens is unlike most other tissues in that it continually grows in size throughout life. This occurs through lifelong proliferation of epithelial cells, and their differentiation into new fiber cells. These newly formed fiber cells add onto existing generations of fiber cells resulting in concentric layers of fiber cells forming a radial gradient of age (Bassnett et al., 1999; Kuszak et al., 2006a; Kuszak et al., 2004a). Newly formed fiber cells reside at the cortical region of the lens while the oldest fiber cells, that formed the lens during embryonic development, are at the core of the lens. Studies show that this continued accumulation of fiber cells results in increasing lens volume and weight with age(Augusteyn, 2008, 2010, 2014a, b).

The relationship between lens size and biomechanical stiffness remains unclear. In support of lens size contributing to lens stiffness, in a comprehensive study of age-related changes in the mouse lens, we found that a nearly linear increase in lens volume parallels a substantial increase in lens stiffness up to 1 year of age (Cheng et al., 2019). However, the lens does not increase in size indefinitely, likely due to space and size limitations within the eye (Augusteyn, 2008). Similarly, as lens volume plateaus with increasing age, the biomechanical stiffening of the lenses also tapers off and is not greatly increased between 18-30 months of age (Cheng et al., 2019), correlating with the plateau in lens growth. Additional support has been provided by other studies (Heys et al., 2004). For example, knockout of a key lens intermediate filament protein CP49 (BFSP2) results in reduced lens size as well as decreased biomechanical stiffness (Fudge et al., 2011). Nevertheless, further studies are required to evaluate this size-stiffness relationship in more detail.

Additional structural features other than lens size likely contribute to lens biomechanical stiffness. In a study on the multiscale transfer of load from the macro-to micro-scale level, we determined that gross lens shape change results in microstructural and cellular changes at the peripheral regions of the lens. At the anterior region of the lens, gross lens shape change induced by compression of lenses results in thinning of the collagenous lens capsule, an increase in lens epithelial cell area, and opening of the Y-shaped mouse suture (Parreno et al., 2018). Additionally, at the equatorial region, lens shape change results in an increase in cortical fiber cell widths. Since our morphometric evaluation of microstructural and cellular changes in aging lenses reveal that these features also change with age as the lenses become stiffer, age-related structural changes such as lens capsule thickening, epithelial cell area expansion, or fiber cell widening (Cheng et al., 2019) may contribute to a stiffer lens. Additionally, others have also suggested that the central core (lens nucleus) is an important determinant of lens biomechanical properties and ability for shape change. The lens nucleus is formed via compaction of fiber cells, resulting in a hardened lens core that is orders of magnitude greater in stiffness than the cortical fiber cells (Heys et al., 2004). In humans, as the lens changes shape during visual accommodation, the lens nucleus also deforms (Brown, 1973; Hermans et al., 2007; Patnaik, 1967). Furthermore, the lens nucleus becomes larger and stiffer with aging in humans and mice (Cheng et al., 2019; Heys et al., 2004), and is thus thought to be responsible for whole lens stiffening with age (Heys et al., 2004).

In this study, we tested the hypothesis that lens size is a determinant of lens stiffness and investigated which structural features in the lens might be correlated with size and stiffness. To test this hypothesis, we used an allometric approach by first comparing the biomechanical properties between three rodent species – mouse, rat and guinea pig. Since lens size scales in proportion to organism size, this approach enabled us to test the relationship of lens size and biomechanical properties while controlling for lens age. We found mouse lenses to be softer than the larger rat or guinea pig lenses in agreement with our initial hypothesis that lens stiffness scales with size. However, we determined rat lenses to be stiffer than guinea pig lenses despite being smaller than guinea pig. Therefore, we next investigated what structural features could account for the elevated stiffness in the rat lenses versus guinea pig lenses. We compared lens microstructural features (capsule thickness, epithelial cell size, fiber widths, nuclear size) in rat versus guinea pig lenses. Our data suggests that the difference in lens biomechanical stiffness between rat and guinea pig may be attributed to differences in the size of the lens nucleus with respect to the size of the lens (i.e., the nuclear fraction of the lens).

## MATERIALS AND METHODS

### Rodent lenses and lens images

Mouse care and euthanasia procedures were approved by the Institutional Animal Care and Use Committee at the University of Delaware. All procedures were conducted in accordance with the Association for Research in Vision and Ophthalmic and Vision Research (ARVO) Statement for the use of Ophthalmic and Vision Research, and the Guide for the Care and Use of Laboratory Animals by the National Institutes of Health.

Mouse eyes were enucleated from wild-type C57BL/6 mice between the ages of 7 and 10 weeks. Long Evans Rat and Guinea pig eyes from animals between the ages of 7 and 10 weeks were purchased (BioChemed Services; Winchester, VA). Eyes were shipped overnight in phosphate buffered saline (PBS) within conical tubes. The eyes were kept cold by surrounding the conical tubes with cooling packs in a sealed Styrofoam container.

Lenses were dissected from eyeballs in PBS (137mM NaCl, 2.7mM KCl, 8.1mM Na_2_HPO_4_, 1.5mM KH_2_PO_4_, pH 8.1) as previously described (Parreno et al., 2018). Briefly, the optic nerve was removed from eyeballs using microdissection scissors, which were then used to cut from the posterior to the anterior region of the eyes. Finally, the lenses were released by applying pressure to the uncut sides of the eye.

### Biomechanical testing of lenses and imaging

The compressive properties of lenses were assessed using load-controlled, sequential application of glass coverslips onto lenses as previously performed (Cheng et al., 2016; Cheng et al., 2019; Gokhin et al., 2012; Parreno et al., 2018) with minor modifications. Briefly, dissected lenses were placed within a bespoke loading chamber (Gokhin et al., 2012). Coverslip loads, with an average weight of 129.3mg, were applied sequentially, two coverslips at a time, onto the lenses. To allow for stress-relaxation equilibration, the lenses were compressed for 2 minutes prior to image acquisition. After 2 minutes, side view images of the lenses were captured, through a 45° angled mirror that was placed at a fixed distance from the lens, on a Nikon Coolpix digital camera connected to an Olympus SZ11 dissection microscope.

After the removal of the terminal 2586mg (20 coverslip) load, the wet weights of lenses were measured using an analytical weighing scale (Mettler-Toledo; Columbus, OH). Finally, the lens hardened nuclear masses (center core region of the lens) were isolated from lenses. This was achieved by removing the lens capsule and dissociating the soft cortical fiber cells by gently rubbing between gloved fingertips. The remaining tissue was the hardened nuclear core. Digital images were then obtained as described above.

### Gross lens morphometric analysis of images

The axial and equatorial diameters of lenses and nuclear regions were measured using FIJI software. Lens strain was calculated using the equation ε = (d-d_0_)/d_0_, where ε is strain, d is axial or equatorial diameter at a given load, and d_0_ is the initial axial or equatorial diameter before the application of any load (0 coverslips). Lens volume was calculated using the equation, lens volume = 4/3 x π x r_E_^2^ x r_A_, where r_E_ is the equatorial radius and r_A_ is the axial radius. Nuclear volume was calculated using the equation nuclear volume = 4/3 x π x r_N_^3^, where r_N_ is the radius of the lens nucleus. Nuclear fraction was calculated using the formula, nuclear fraction = nuclear volume/lens volume.

### Whole mount imaging preparation of fixed lenses

Whole mount imaging was performed as previously described (Parreno et al., 2018). Freshly dissected lenses were fixed in 4% paraformaldehyde in PBS at room temperature. After 30 minutes, lenses were washed three times (5 minutes per wash) in PBS and then incubated in permeabilization/blocking solution (PBS containing 0.3% Triton, 0.3% bovine serum albumin, and 3% goat serum) at room temperature for another 30 minutes. Next, lenses were placed in staining solution containing fluorescent CF640 dye conjugated to wheat-germ agglutinin (Biotium, Fremont, CA) (1:500), Hoechst 33342 (Biotium) (1:500), and rhodamine-phalloidin (Thermo Fisher Scientific) (1:20). After an overnight incubation, lenses were washed three times in PBS for 5 minutes before performing confocal microscopy.

### Confocal microscopy

Confocal microscopy on lens whole mounts were performed on a Zeiss LSM880 laser-scanning confocal fluorescence microscope (Zeiss, Germany) as previously described (Parreno et al., 2018). To image the anterior capsule, epithelial cells and suture, lenses were placed anterior side down on 10mm microwell glass-bottomed dishes (MakTek, Ashland, MA). To prevent lens movement while imaging, the lenses were immobilized within a small circular divot that was created in a thin layer of 2% agarose in PBS using a 2mm biopsy punch (Accu-punch, Acuderm inc., Fort Lauderdale, FL). To image the posterior suture, lenses were placed posterior side down within the agarose divot. Z-stack images were acquired with a 40x oil Plan-Apo 1.4 NA objective using a step size of 0.3μm. To image the equatorial fiber cells, the lenses were placed in optical glass bottomed Fluorodishes (World Precision Instruments, Sarasota, FL) and balanced on the side using agarose wedges (Cheng et al., 2017; Parreno et al., 2018). Z-stack images were acquired with a 20x air 0.8NA objective using a step size of 0.5μm.

### Analysis of microstructural and cellular features

Raw fluorescent images were processed using Zen Black 2.3SP1 (Zeiss) software. FIJI software was used for lens morphometric analysis and measurements of microstructural features as previously described (Parreno et al., 2018). Capsule thickness was measured by obtaining intensity distributions of capsular (WGA-640) and basal epithelial F-actin (rhodamine-phalloidin) stains using line scan analysis of XZ plane-view reconstructions and by performing subtractive peak-to-peak analysis of fluorescent pixel intensity to obtain distance (Parreno et al., 2018).

Epithelial cell area was calculated by tracing a population of at least 30 cells (region of interest; ROI) whose boundaries were identified by staining of F-actin, using rhodamine-phalloidin, at cell membranes. The total number of cells within the ROI was determined by counting cell nuclei that were stained with Hoechst. Average cell number was calculated using the equation, average cell area = ROI area / total number of cells.

Fiber cell width was calculated by line scan analysis of fiber cell membranes stained with rhodamine-phalloidin at the lens equator, ∼10μm inward from the fulcrum. On FIJI, the Distributed Deconvolution (Ddecon) plugin with Z-line predictive model was used to provide high spatial precision when analyzing fiber cell widths.

### Statistical Analysis

Each experiment was replicated at least four times. The sample size (N) of each experiment is indicated in figures. Differences between multiple groups of data were assessed using a one-way analysis of variance followed by a Tukey’s multiple comparisons post hoc test. Differences between two groups of data were detected using unpaired *t* tests.

## RESULTS

### Lens size scales with overall rodent size

To characterize the allometric properties of rodent lenses, we used 7–10-week-old mice, rats, and guinea pigs. The mice are the smallest of the rodents weighing 28.1 ± 0.6g (Figure 1A; average ± S.E.). Rats have an average weight of 252.2 ± 25.0g which is 9.0-fold more than mouse. Guinea pigs weigh the most of the three species weighing 571.3 ± 33.9g, which is 20.3-fold more than mouse and 2.3-fold more than rats. Next, we dissected the lenses (Figure 1B) and measured their wet weight (Figure 1C). Mouse lenses weigh the least, with an average wet weight of 5.8 ± 0.01mg. Rat lenses have a wet weight of 43.3 ± 4.7mg, which is approximately 7.5-fold greater than mouse lenses. Guinea pig lenses are the heaviest of the three with an average wet weight of 74.7 ± 1.5mg, which is 12.8-fold heavier than mouse lenses and 1.7-fold heavier than rat lenses.

**Figure 1.**
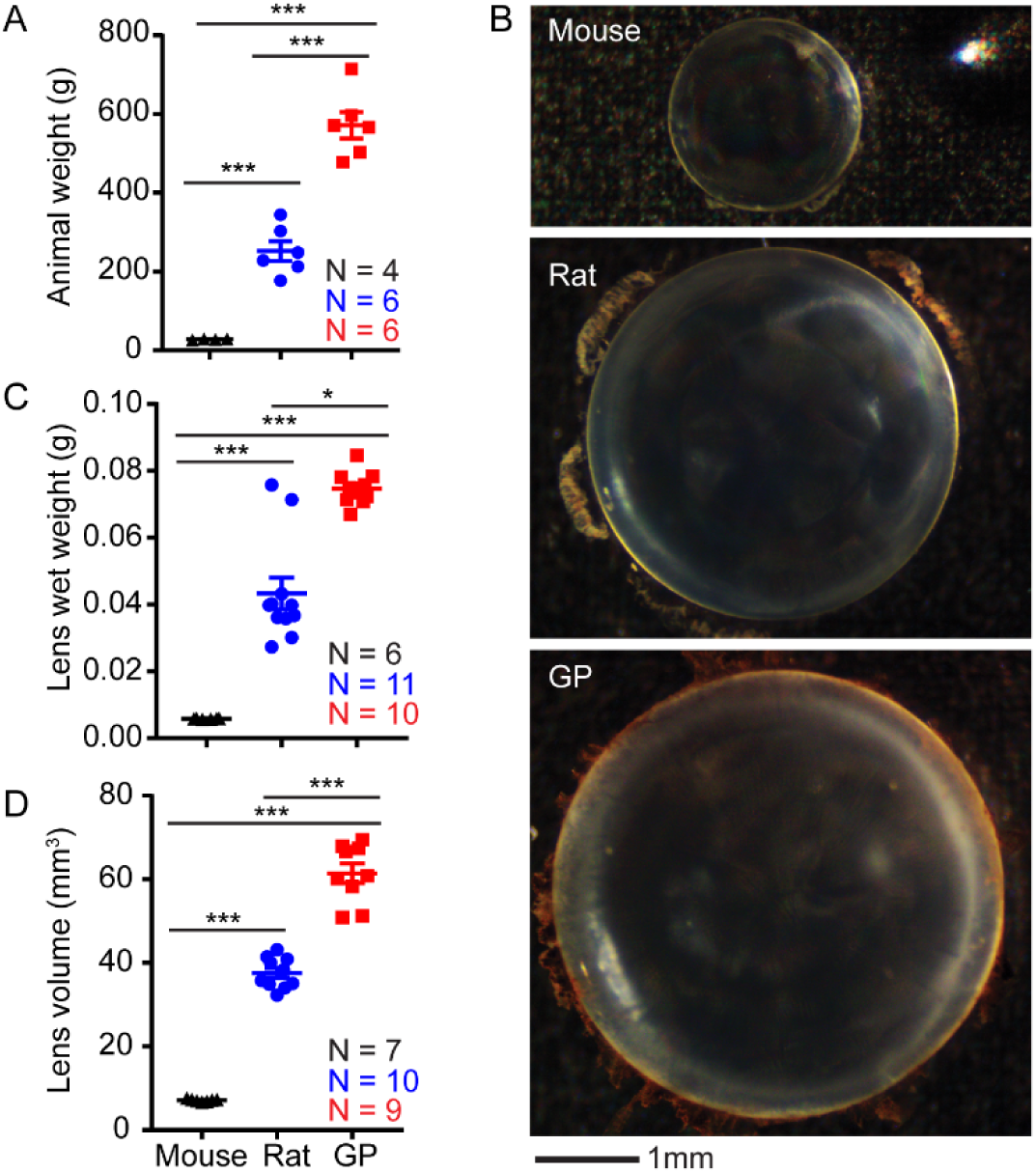
Lens size scales with rodent size. (A) Body weight difference of mouse, rats, and guinea pigs parallel the differences seen in lens size: (B) Top view images of lenses, (C) measured wet weights of lenses, and (D) calculated gross volumes of lenses from mouse, rat, and guinea pig (GP). The proportion of lens to animal weights is comparable between species (0.020% for mouse, 0.017% for rat, and 0.013% for guinea pig). *, p < 0.05; ***, p < 0.001

Finally, we measured the axial and equatorial diameter of lenses and calculated lens volume (Figure 1D). Mouse lenses are the smallest of the three species with an average volume of 7.1 ± 0.1mm^3^. Rat lenses have an average volume of 37.6 ± 1.2mm^3^ which is 5.3-fold greater than mouse. Guinea pig lenses are the largest of the three species with an average volume of 61.5 ± 2.3mm^3^, which is 8.6-fold larger than mouse lenses and 1.6-fold larger than rat lenses.

### Lens biomechanical stiffness does not necessarily scale with lens size

To examine if lens biomechanical stiffness scales with lens size, we performed load-controlled (coverslip) compression of lenses followed by calculation of axial compressive strain (negative) and equatorial expansion strain (positive). Based on their larger size, we hypothesized guinea pig lenses would be the stiffest of the three species, followed by rat then mouse. As expected, mouse lenses are the softest of the three rodent lenses, demonstrated by the highest absolute axial and equatorial strains (Figure 2A-C). However, when we compared rat versus guinea pig biomechanical properties, we determined that lens axial strain is similar between guinea pig and rat lenses (p > 0.05; Figure 2B and 2D). We also found that guinea pig lenses underwent greater equatorial expansion when loaded with 1552-2327mg (12-18 coverslip) loads as compared to rat lenses (Figure 2C and 2E). The greatest difference is seen at a load of 1810mg (14 coverslips) where equatorial strain is 4.7 ± 0.3% for rat lenses and 6.7 ± 0.8% for guinea pig lenses. While axial strain at 14 coverslips is not statistically different in rat versus guinea pig lenses, the axial strain is −13.5 ± 0.7% for rat lenses and −15.1 ± 0.8% in guinea pig lenses with p values approaching significance (p = 0.08; Figure 2D).

**Figure 2.**
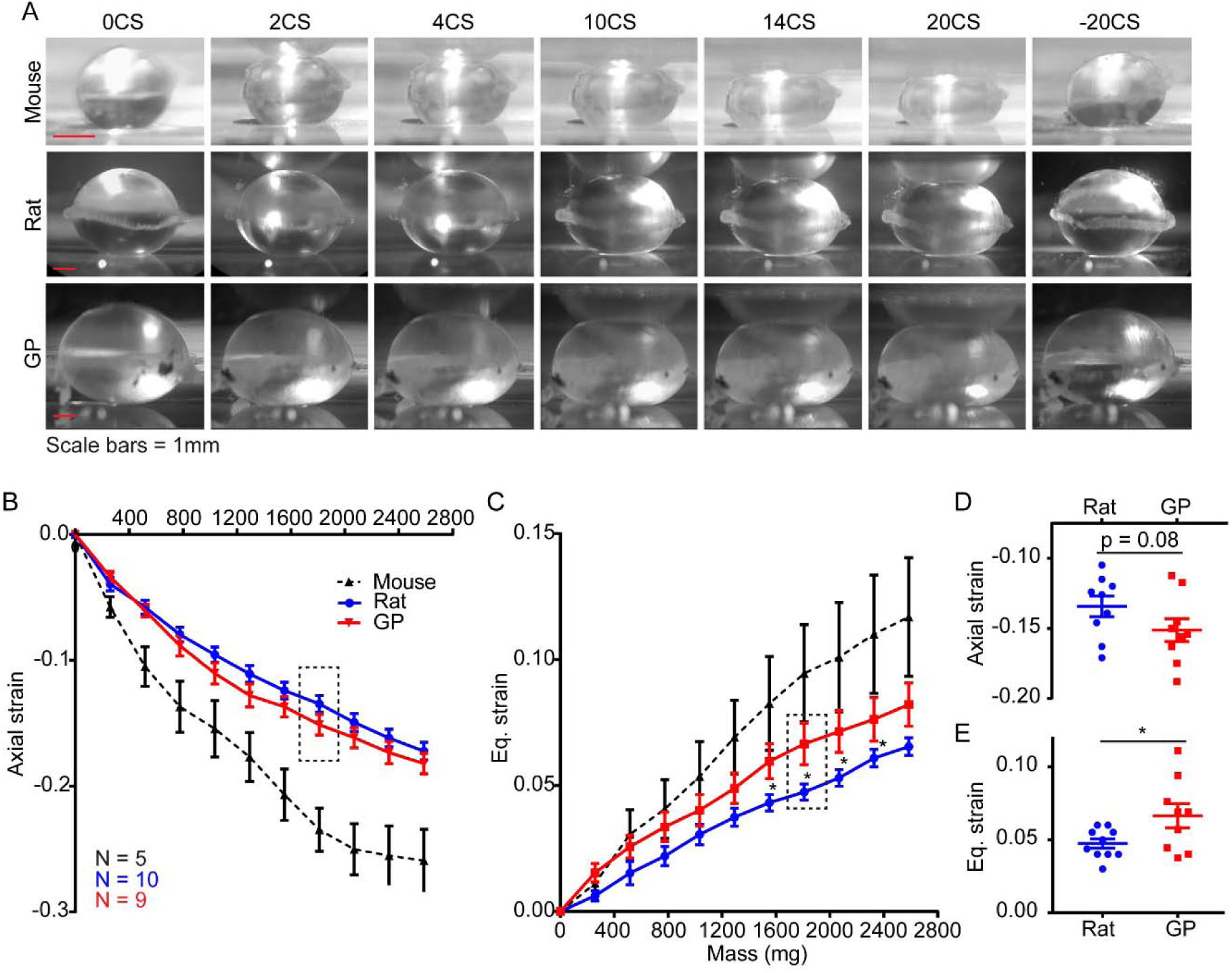
Differences in lens biomechanical properties. (A) Side view images of mouse, rat, and guinea pig (GP) lenses. Mouse lenses are overall softer than both rat and guinea pig lenses as determined by measuring (B) axial and (C) equatorial (eq.) strain of lenses. While guinea pig lenses are larger than rats, they were not stiffer. (B) Axial strain was similar between rat and guinea pigs at all compressive loads tested, although at a load of (D) 1810.2mg (14 coverslips), there was a non-significant trend toward higher absolute axial strain in guinea pig lenses (p = 0.08). (C) Equatorial strain measurements revealed guinea pig lenses are softer between 1551.6-2327.4mg (12-18 coverslip) loads. These differences in equatorial strain were most pronounced at a load of (E) 1810.2mg (14 coverslip). *, p < 0.05 between rat and guinea pig.

As we expected, rat and guinea pig lenses are stiffer than mouse lenses. However, we were surprised to find that although rat lenses are smaller than guinea pig lenses, they are stiffer. This indicates that lens biomechanical stiffness does not necessarily scale with lens size. Since we found previously that certain microstructural (capsule thickness) and cellular (epithelial cell area, fiber widths, nuclear size) features increase with age as the lens continues to grow in size, we questioned whether differences in these microstructural/cellular features could account for stiffer rat lenses in comparison to guinea pig lenses. Therefore, we compared the microstructural and cellular features between rat and guinea pig lenses.

### Lens microstructural features — capsule thickness, epithelial cell area, cortical fiber cell widths— all scale with the lens size for rats and guinea pigs

We performed an analysis of lens capsule and cell sizes in rat and guinea pig lenses. Whole lenses were fixed and incubated in WGA and Phalloidin, which stains the lens capsule matrix(Parreno et al., 2018) and the underlying F-actin at the epithelial cell membranes, respectively (Figure 3A). This allows us to measure capsule thickness by performing line scan analysis on sagittal (XZ plane) optical sections of reconstructed images (Cheng et al., 2019; Parreno et al., 2018). We determined that rat lenses have an average capsule thickness of 8.1 ± 0.7μm (Figure 3B), while guinea pigs have an average lens capsule thickness of 14.8 ± 0.9μm, which is 1.8-fold thicker than rat lens capsule.

**Figure 3.**
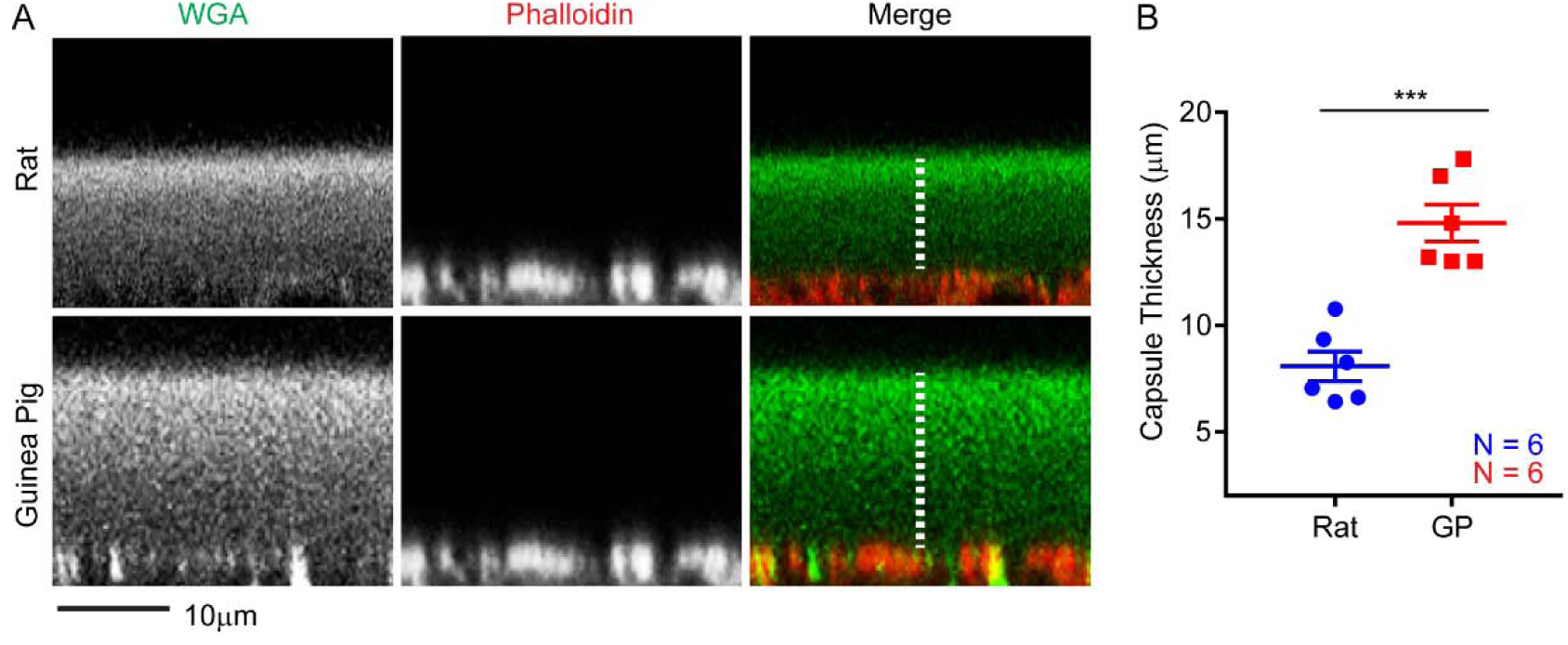
Lens capsule thickness measurements. (A) representative (XZ reconstruction) confocal images and (B) measured thicknesses of capsules from rat and guinea pig lenses. Lenses were stained with WGA (green) to visualize lens capsule, and phalloidin (red) to visualize basal F-actin in epithelial cells at the epithelial-capsule interface. ***, p < 0.001

Underlying the capsule at the anterior region of the lens are epithelial cells. Staining lenses with phalloidin allows us to visualize F-actin associated with cell membranes at the lateral boundaries of the anterior epithelial cells (Figure 4A), enabling us to measure epithelial cell area. We found that rat lenses have an average epithelial cell area of 166.1 ± 19.0μm^2^ (Figure 4B), whereas guinea pigs have an average lens epithelial cell area of 346.7 ± 11.5μm^2^, which is 2.1-fold larger than rat lens epithelial cells.

**Figure 4.**
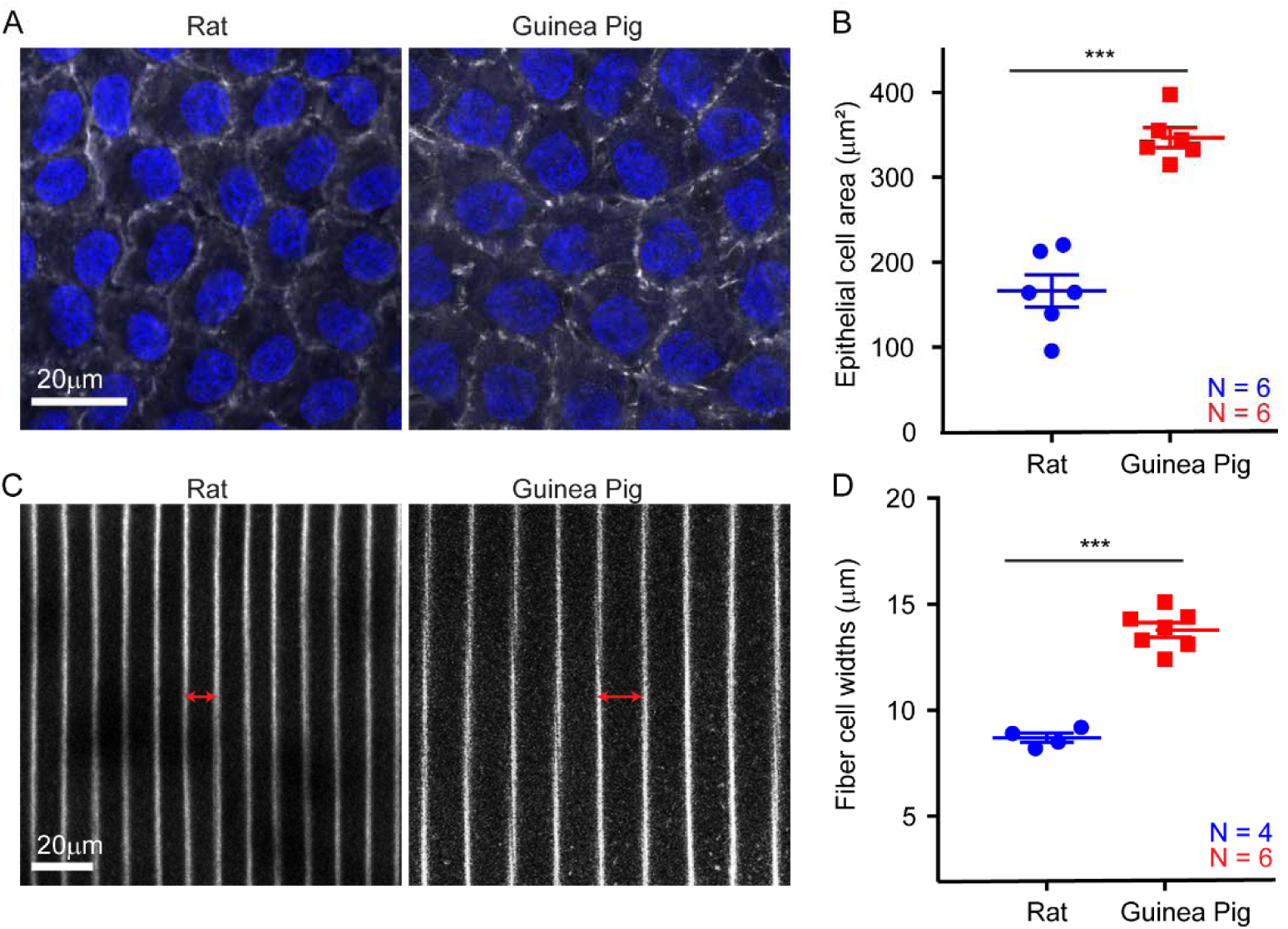
Lens epithelial cell and fiber cell widths are greater in guinea pig lenses. (A) Representative confocal images of rat and guinea pig anterior epithelial cells; and (B) corresponding epithelial cell area measurements demonstrating larger epithelial cells in guinea pig than rat lenses. (C) Representative equatorial fiber cell widths; and (D) corresponding fiber cell width measurements demonstrating wider fiber cells in guinea pig than rat lenses. Lenses were stained with phalloidin (gray scale) to visualize F-actin at cell boundaries, and Hoechst (blue) to visualize epithelial cell nuclei. ***, p < 0.001

Next, we measured the cortical fiber cell widths at the equator in rat and guinea pig lenses (Figure 4C). We observed that rat lenses have an average fiber cell width of 8.7 ± 0.2μm (Figure 4D), whereas guinea pigs have an average fiber cell width of 13.8 ± 0.3μm, which is 1.6-fold greater than rat fiber cell widths. Thus, the capsule thickness, epithelial cell area, cortical fiber cell widths all scale with the lens size for rats and guinea pigs.

Finally, since a high level of suture branching as seen in primate lenses may be indicative of lens flexibility and propensity for lens shape change (Kuszak et al., 1984; Kuszak et al., 2006a; Kuszak et al., 2004b; Sivak et al., 1994), we examined whether guinea pig suture branching differed from rat. The anterior and posterior suture organization in both rat and guinea pig lenses are similar with only 3-4 suture branches at both the anterior and posterior regions of the lens (Figure 5).

**Figure 5.**
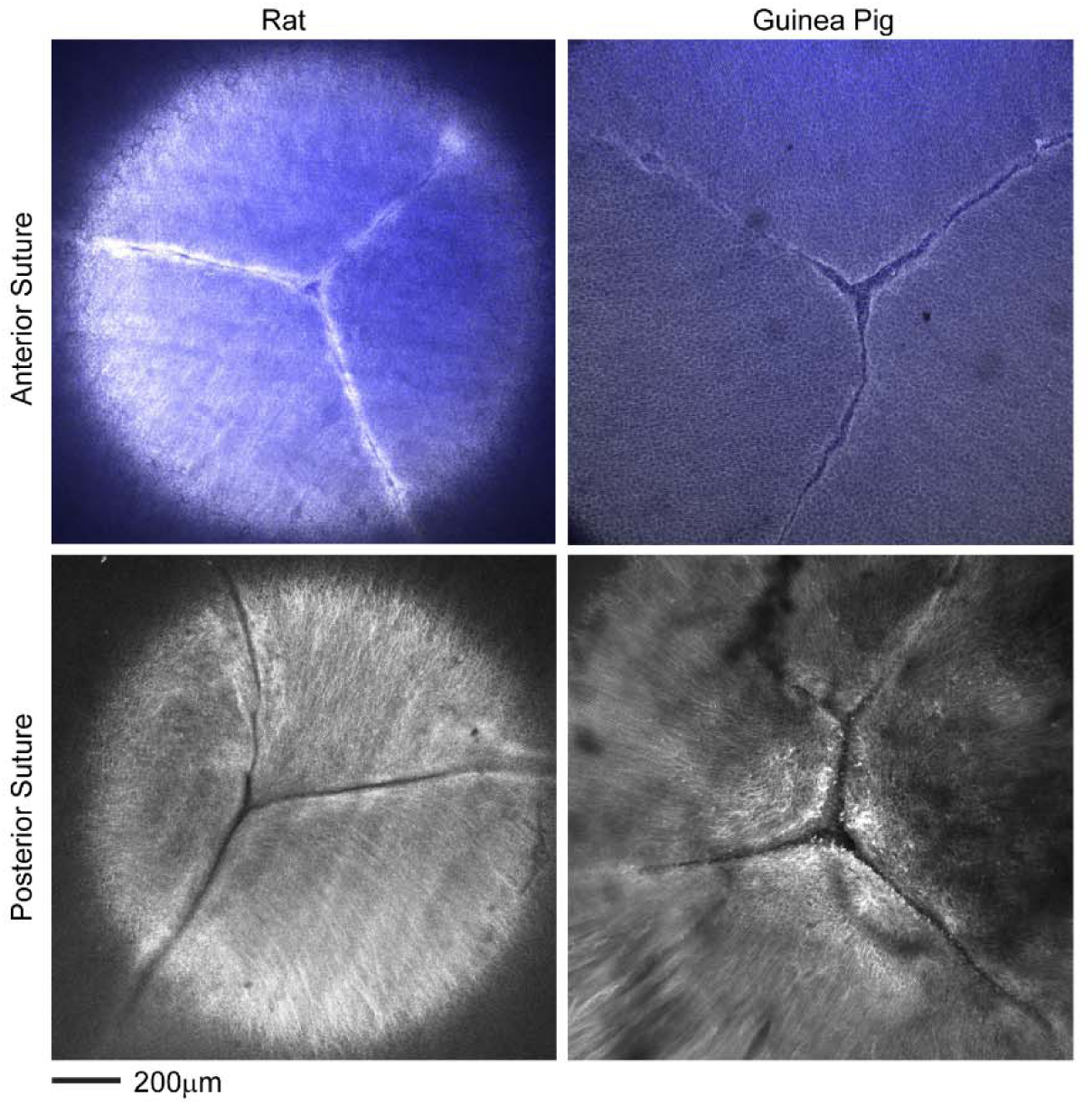
Suture organization in rat and guinea pig lenses. Lenses were stained with phalloidin (gray scale) to stain F-actin at cell boundaries, and Hoechst (blue) to stain nuclei (only present at the anterior region of the lens). This revealed Y-shaped sutures at the anterior (top panels) and posterior (bottom panels) regions of the lenses.

### Nuclear fraction does not scale with lens size for rat and guinea pig

Fiber cells at the innermost, center (nuclear) regions of the lens become compacted (Al-Ghoul et al., 2001; Heys et al., 2004) forming a hardened spherical nuclear structure in mouse lenses (Cheng et al., 2018; Cheng et al., 2019; Gokhin et al., 2012). There is a marked increase in size and stiffness of the nucleus in both mouse and human lenses with aging (Cheng et al., 2019; Heys et al., 2004). Therefore, we next explored whether nuclear size was different between rat and guinea pig lenses. To investigate nuclear size, we calculated nuclear volume by measuring the nuclear axial and equatorial diameters. We determined that despite the difference in whole lens volume (Figure 1), rat and guinea lenses have similar nuclear volumes (p = 0.62). On average, rats have a nuclear volume of 12.5 ± 1.5mm^3^ and guinea pig lenses have an average nuclear volume of 13.8±2.0mm^3^. We next calculated the nuclear to lens volume ratios. We determined that the rat lens nuclear fraction is 29.4 ± 3.5% of the entire lens, whereas the guinea pig lens nuclear fraction occupies a significantly smaller fraction of the total lens volume at 20.0 ± 2.6% (Figure 6).

**Figure 6.**
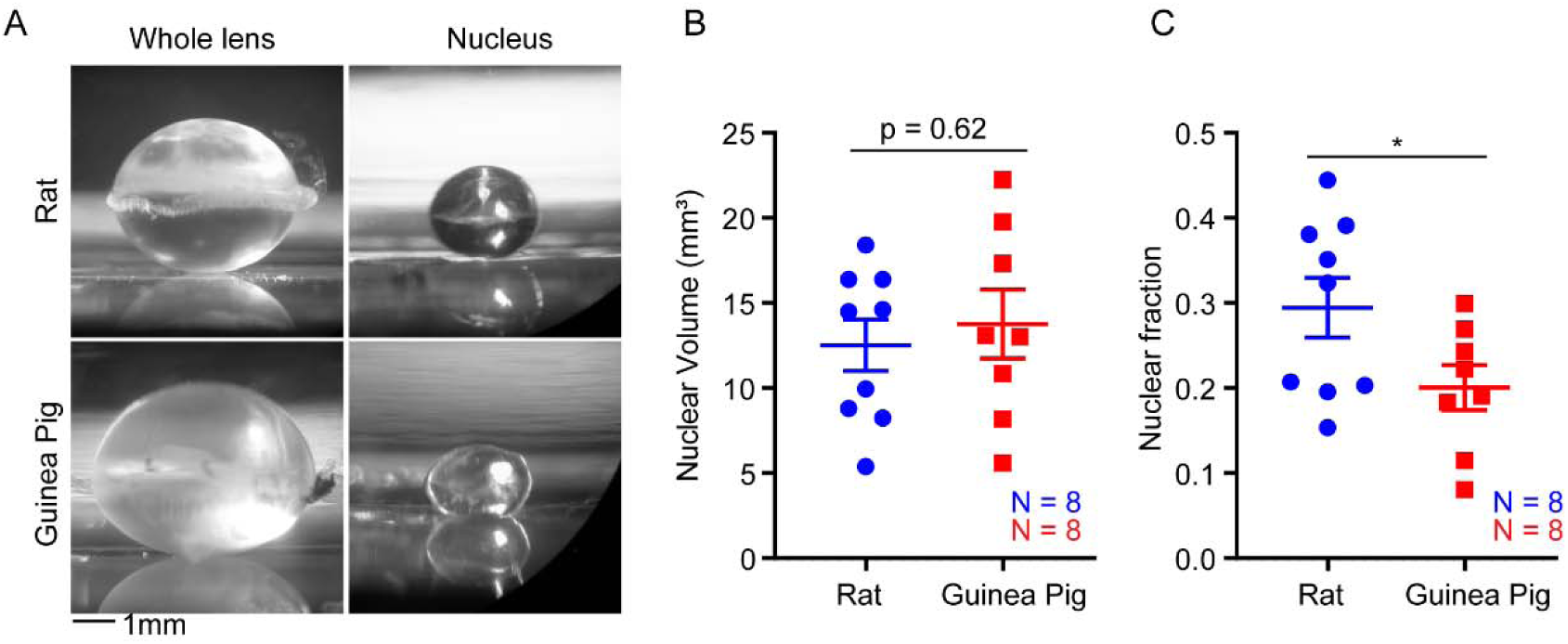
Lens nuclear fraction is larger in rat than guinea pigs. (A) Representative side view images of whole lenses (left panels) and hardened nuclear regions of rat and guinea pig lenses (right panels). (B) Rat and guinea pig nuclear regions were similar in volume but occupied a greater (C) proportion of the lens (i.e., nuclear fraction of the lens). *, p < 0.05

## DISCUSSION

Guinea pigs have larger lenses than rats; however, we show that these lenses are biomechanically softer. Therefore, lens size is not always predictive of lens biomechanical stiffness. Based on this finding, the question arises: what lens features make rat lenses stiffer than guinea pig? To permit gross lens shape change, it has been shown that capsular, cellular, and nuclear deformation is required (Brown, 1973; Hermans et al., 2007; Parreno et al., 2018; Patnaik, 1967; Weeber et al., 2005). It is also known that as lenses age and become stiffer, these features also change with age including lens capsule thickening, epithelial cell area expansion, fiber cell widening, and nuclear enlargement (Balaram et al., 2000; Cheng et al., 2019; Danysh et al., 2008). However, it remains unclear how differences in these structures may relate to lens stiffness. Therefore, we explored the possibility that the higher biomechanical stiffness of rat lenses is due to capsular, cellular, and/or nuclear features that may not scale with lens size. To our knowledge we are the first to comprehensively examine capsular thickness, epithelial cell area, fiber cell widths, and nuclear sizes in guinea pig lenses in comparison to those of rats.

Capsular thickness does not explain the stiffer lens phenotype in rat as compared to guinea pig. Since a decrease in capsule thickness accompanies whole lens shape change (Parreno et al., 2018), we investigated the possibility that rats may have thicker lens capsules than guinea pig, thereby impeding lens shape change. However, our findings show that capsular thickness scales with lens size - guinea pigs have thicker capsules than rat – so that this does not explain the stiffer rat lens compared to the guinea pig.

Next, we examined lens epithelial cell area as a determinant of lens stiffness. We have shown that epithelial expansion is required for lens shape change(Parreno et al., 2018). This leads to a postulation that larger, flatter epithelial cells do not have the capability to expand as much as smaller epithelial cells, potentially restricting whole lens shape change as would be indicated by stiffer lenses. Thus, we explored the possibility that epithelial cell areas do not scale with lens size. However, our findings show that lens epithelial cell area scales with lens size, as guinea pig epithelial cells are larger than rat. We conclude that epithelial cell area does not explain the increased biomechanical stiffness in rats as compared to guinea pigs.

In addition to epithelial cell area, we examined fiber cell widths as a determinant of lens stiffness. Since fiber cell widening accompanies lens shape change(Parreno et al., 2018) it may be postulated that wider cells have less ability to expand than thinner cells during lens shape change, therefore we considered whether fiber cell widths might be larger in the stiffer rat lens as compared to the softer guinea pig lenses. However, our findings reveal that fiber cell widths scale with lens size but not stiffness and are larger in guinea pig than rat. Therefore, expanded fiber cell widths cannot explain the elevated biomechanical stiffness of rat lenses.

The increase in lens stiffness during aging coincides with an increase in lens capsule thickness, increase in epithelial cell area, and fiber cell widths (Balaram et al., 2000; Cheng et al., 2019; Danysh et al., 2008; Fisher and Pettet, 1972; Hara and Hara, 1988; Krag et al., 1997) as characterized in mouse and/or human lenses. Our allometric comparison between rat and guinea pig lenses demonstrates that these features do not determine lens biomechanical properties. Further support that capsule thickness, epithelial cell area, and fiber cell widths do not determine lens biomechanical properties is provided by comparing our current findings for these rat lens features with mouse lens features, which we characterized previously (Cheng et al., 2019; Parreno et al., 2018). Despite mouse lenses being substantially softer than rat (Fig. 2), we find that mouse lens capsules are thicker, epithelial cell areas are equivalent between mouse and rat lenses, and equatorial fiber cell widths are greater in mouse than rat lenses (Table S1) (Parreno et al., 2018). These comparisons between mouse and rat lenses not only demonstrate that these microstructural features do not determine differences in lens biomechanical stiffness for mouse and rat, but also show they do not necessarily scale with lens size.

In the present study, we also considered the possibility that the structural organization of the lens suture in rat and guinea pig could vary and lead to differences in biomechanical lens stiffness. Mouse and rat lenses with limited shape change ability have 3 to 4 suture branches (Parreno et al., 2018), with the 3 branch sutures often described as being Y-shaped (Kuszak et al., 2004b; Sivak et al., 1994). In contrast to rodent lenses, primate lenses that have the capacity to undergo significant lens shape change have multiple suture branches and are described as being star-shaped (Kuszak et al., 2006b). This multi-branched suture organization has been suggested to influence the ability of lenses to change shape and accommodate (Kuszak et al., 2006a). To our knowledge the suture organization of guinea pigs had not been studied. Here, we determined that guinea pig lens sutures have 3-4 suture branches and are Y-shaped, similar to those of the rat lens. Thus, differences in suture organization between rat and guinea pig do not account for the increased stiffness of rat lenses.

While the aforementioned features cannot explain the increased stiffness of rat versus guinea pig lenses, the lens nucleus may determine whole lens biomechanical properties. This idea is suggested by observations that increased lens nuclear stiffness and nuclear size parallels elevated whole lens stiffness with age (Cheng et al., 2019; Heys et al., 2004). Therefore, we tested the possibility that lens nuclear size may account for elevated biomechanical stiffness of rat lenses in comparison to guinea pig. We determined that despite guinea pig lenses being 1.6-fold greater in volume than rat lenses, the nuclear volume was similar between guinea pigs and rats, so that the proportion of the lens occupied by the nucleus in the guinea pig lens was smaller than in the rat. The lens nucleus is markedly stiffer than cortical fiber cells and is considered to play a large role in the ability for whole lens shape change (Augusteyn, 2018; Brown, 1973; Hermans et al., 2007; Heys et al., 2004; Patnaik, 1967). Out of all the structural features examined, the fraction of the lens occupied by the nuclear portion is the only feature we determined not to scale with lens size, so that the larger nuclear fraction in rat as compared to guinea pig may explain the biomechanical differences in lens stiffness between rat and guinea pig. In support of this, mouse lenses which are softer than both guinea pig and rat lenses have a nuclear fraction of 11.0±0.2% (Table S1), which is a smaller fraction than either rat (29.5±3.5%) or guinea pig lenses (20.0±2.6%). Therefore, we conclude that the higher lens stiffness in rat versus guinea pig lenses is attributed to a larger nuclear fraction impeding the ability for rat lens shape change.

It is important to note that lens stiffness is complex and not necessarily determined by one structural feature. While our findings suggests that the lens nucleus is a determinant of lens stiffness, findings from our other studies demonstrate that it is not the only feature that contributes to lens stiffness. Previously, we have shown a dramatic increase in nuclear size between 24 and 30 months of age in mice, however, the overall biomechanical stiffness of lenses does not increase over this time period (Cheng et al., 2019). Other factors, such as cytoskeletal networks, are known to contribute to lens stiffness. We found that reduction of the F-actin binding protein, Tropomyosin 3.5, results in softer lenses. This is despite a greater nuclear fraction in Tropomyosin 3.5-depleted mouse lenses (Cheng et al., 2018). Additional support that F-actin networks regulate lens stiffness has also been provided by disruption of the F-actin adaptor molecule, ankyrin-B (Bennett and Healy, 2009). Ankyrin-B links spectrin-F-actin networks to cellular membranes and ankyrin-B knockout lenses have reduced lens stiffness (Maddala et al., 2016). Additionally, intermediate filament proteins also regulate lens biomechanical properties as knockout of specialized intermediate filament protein CP49 (phakinin) reduces lens stiffness (Fudge et al., 2011). The F-actin and intermediate filament networks are biomechanically linked and converge functionally. We have shown that CP49 synergizes with F-actin pointed end capping protein, Tmod1, to control lens stiffness, since deletion of both CP49 and Tmod1 further reduces lens biomechanical stiffness as compared to CP49 deletion alone (Gokhin et al., 2012). The lens contains multi-scale structural elements, and disruption of molecular scale cytoskeletal proteins lead to whole organ level (lens) biomechanical consequences.

Additional biophysical and biochemical factors may also regulate lens stiffness. In addition to a smaller nuclear fraction, guinea pig lenses are reported to have soft nuclei in comparison to rat (Ozaki and Mizuno, 1992) which could also account for the greater whole lens stiffness in rat versus guinea pig lenses. The biophysical properties of other lens microstructures, such as the lens capsule, could also determine lens stiffness. Since lens capsular stiffness also increases with age (Krag et al., 1997), the molecular composition and organization of the capsule may contribute to whole lens stiffening, and will require further investigation. In addition, the biochemical composition of the lens fiber cell bulk may play a role in lens stiffness. Guinea pig lenses contain zeta-crystallins, which are only expressed in a limited number of animal species (Garland et al., 1991; Huang et al., 1987). Lenses with zeta-crystallins have a lower refractive index which may be indicative of reduced protein aggregation as compared to lenses that predominantly contain beta-crystallins (de Castro et al., 2020; Muir et al., 2020; Tardieu et al., 1992; Zhao et al., 2011). Additionally, alpha crystallins could also play a role in determining lens stiffness. In young humans, the nucleus contains predominantly alpha crystallins, a heat-shock protein which tightly binds water (Babizhayev et al., 2003; Babizhayev et al., 2002). However, the proportion of alpha-crystallins in the nucleus decreases with age resulting in aggregation of other crystallins, nuclear dehydration and increased lens nuclear stiffness (Heys et al., 2004; Heys et al., 2007; McFall-Ngai et al., 1985; Ozaki and Mizuno, 1992; Roy and Spector, 1976). Crystallin aggregation and protein cross-linking play a role in lens biomechanics and may be a target against presbyopia (Garner and Garner, 2016; Heys et al., 2007; Nandi et al., 2020; Randall and Vaughan, 1982; Soergel et al., 1999). However, crystallins can also interact with F-actin and intermediate filament networks of the lens (Andley, 2009; Clark et al., 1999; Wang and Spector, 1996). Therefore, the regulation of lens biomechanical properties is likely regulated by multiple factors that interact.

The lens is a unique tissue. In this study we examined the biomechanical properties of three rodent species. It is important to note that in comparison to humans, the visual system of rodents has been regarded as relatively less developed. Rodents have limited visual acuity relying on other senses such as olfactory, tactile and auditory system for their daily lives (Huberman and Niell, 2011; Peirson et al., 2018; Prusky and Douglas, 2004; Prusky et al., 2000). Since nocturnal rodents, such as mice and rats, function under low light conditions, there would be relatively little need to accommodate (i.e., change lens shape to fine focus light onto the retina). In contrast, diurnal rodents, such as guinea pigs, are crepuscular being most active at dawn and dusk. Therefore, guinea pigs would likely benefit from lens accommodation. Indeed, guinea pigs have been shown to have accommodative capabilities (Ostrin et al., 2014). Since guinea pig lenses can accommodate, the finding that guinea pigs have evolved mechanisms to allow for biomechanically softer lenses than rats may not be completely surprising.

## Acknowledgements

We thank Dr. Melinda Duncan (University of Delaware), Dr. Catherine Cheng (Indiana University School of Optometry), and Dr. Christine Wildsoet (Berkeley School of Optometry) for helpful discussions pertaining to this work.

---

**Table S1.** Summary of lens morphological features from mouse, rat, and guinea pig. Mouse data is from Parreno et al. (2018)(Parreno et al., 2018).

|  | **Mouse** | **Rat** | **Guinea Pig** |
| --- | --- | --- | --- |
| **Lens Volume** | 7.1±0.1 mm<sup>3</sup> | 37.6±1.2 mm<sup>3</sup> | 61.5±2.3 mm<sup>3</sup> |
| **Capsule Thickness** | 12.0±0.7 µm | 8.1±0.7 µm | 14.8±0.9 µm |
| **Epithelial Cell Area** | 165.5±10.6 µm<sup>2</sup> | 166.1±19.0 µm<sup>2</sup> | 346.7±11.5 µm<sup>2</sup> |
| **Fiber Widths** | 11.0±0.1 µm | 8.7±0.2 µm | 13.8±0.3 µm |
| **Nuclear Volume** | 0.4±0.1 mm<sup>3</sup> | 12.5±1.5 mm<sup>3</sup> | 13.8±2.0 mm<sup>3</sup> |
| **Nuclear Fraction** | 11±0.2% | 29.4±3.5% | 20.0±2.6% |

